# A recombinant receptor-binding domain in trimeric form generates completely protective immunity against SARS-CoV-2 infection in nonhuman primates

**DOI:** 10.1101/2021.03.30.437647

**Authors:** Limin Yang, Deyu Tian, Jian-bao Han, Wenhui Fan, Yuan Zhang, Yunlong Li, Wenqiang Sun, Yanqiu Wei, Xiaodong Tian, Dan-dan Yu, Xiao-li Feng, Gong Cheng, Yong-tang Zheng, Yuhai Bi, Wenjun Liu

**Author notes:** These authors contributed equally to this work (Limin Yang, Deyu Tian, Jian-bao Han). Corresponding authors. (W.L.); (L.Y.); (Y.B.); (G.C.); (Y.Z.).

## Abstract

Safe and effective vaccination is critical to combatting the COVID-19 pandemic. Here, we developed a trimeric SARS-CoV-2 receptor-binding domain (RBD) subunit vaccine candidate that simulates the natural structure of the spike (S) trimer glycoprotein. Immunization with RBD-trimer induced robust humoral and cellular immune responses and a high level of neutralizing antibodies that were maintained for at least 4 months. Moreover, the antibodies that were produced in response to the vaccine effectively neutralized the SARS-CoV-2 501Y.V2 variant. Of note, when the titers of the antibodies dropped to a sufficiently low level, only one boost quickly activated the anamnestic immune response, resulting in complete protection against the SARS-CoV-2 challenge in *rhesus macaques* without typical histopathological changes or viral replication in the lungs and other respiratory tissues. Our results indicated that immunization with SARS-CoV-2 RBD-trimer could raise long-term and broad immunity protection in nonhuman primates, thereby offering an optimal vaccination strategy against COVID-19.

## Introduction

Since severe acute respiratory syndrome coronavirus 2 (SARS-CoV-2) was identified as the causative agent for the current coronavirus disease (COVID-19) ^[1]^, the ongoing global pandemic has infected more than 1.6% (over 120 million people) of the world’s population, which has caused a severe economic burden and hindered social development across the globe. There is an urgent need to develop safer and more effective vaccines to end the current health crisis. Researchers worldwide are racing to develop COVID-19 vaccines; more than 200 vaccine candidates are in development, and 70 are in clinical trials. Multiple vaccine development strategies have been pursued simultaneously, including viral vectors^[2–4]^, inactivated whole viruses^[5, 6]^, DNA^[7, 8]^, RNA^[9–11]^, subunits^[12, 13]^, virus-like particles^[14]^, and live attenuated viruses (World Health Organization, WHO). Vaccines with different development strategies have distinct advantages and application limitations. Therefore, the development of new vaccine strategies to avoid shortcomings is still the focus of vaccine researchers.

SARS-CoV-2 is a member of the *Betacoronavirus* genus of the *Coronaviridae* family of enveloped RNA viruses homologous to SARS and Middle East respiratory syndrome (MERS) coronaviruses. The viral surface spike (S) glycoprotein mediates receptor binding and cell entry and is the primary target for vaccine design^[15]^. To mimic the native S trimer structure, an artificially designed S trimer was constructed by fusing the C-terminal region of human type Iα collagen, and this construct was used for vaccine research^[16]^. Previous studies have found that both SARS-CoV and MERS-CoV display antibody-dependent enhancement (ADE), leading to a potential risk of vaccine enhancement of disease^[17, 18]^. The principle is that the nonneutralizing antibodies produced in response to the vaccine mediate virus infection via the Fcγ receptor^[18]^. Considering the high degree of homology of SARS-CoV-2, the potential ADE effect needs to be considered seriously in designing COVID-19 vaccines. To mitigate the ADE effect, minimizing nonneutralizing epitopes and keeping only the critical neutralizing epitope to elicit robust protective immunity is a solution^[19]^. The receptor-binding domain (RBD) located at the C-terminus of the S1 subunit has thus attracted attention. Several lines of evidence have indicated that RBD-specific antibodies could minimize the ADE effect^[20]^, and most discovered potent neutralizing antibody (NAb) target the RBD region^[21, 22]^. SARS-CoV-2 initiates viral replication by binding via the RBD to the cell surface receptor angiotensin-converting enzyme 2 (ACE2). Hence, the RBD can be used as a vaccine target to block viral attachment. Additionally, the RBD possesses T cell epitopes, which can provoke antiviral T cell responses^[11, 12]^. However, the molecular weight of the RBD is small, which leads to its weak immunogenicity. Some researchers have tried to express it by fusion with the Fc domain and prepare it as a dimer, thereby greatly enhancing its immunogenicity^[13]^. Some evidence has shown that the NAb level in SARS-CoV-2-infected humans significantly declines from the second month and can even be lost ^[23]^, allowing them to be reinfected ^[24]^. This short-lasting antibody protection poses a severe challenge for vaccine development. In addition, SARS-CoV-2 is characterized by a high mutation rate, and some evidence has proven that the 501Y.V2 variant (B.1.351) that emerged in South Africa severely reduced the protective efficacy of licensed mRNA vaccines ^[25–27]^. Therefore, a vaccine strategy that can produce persistent and broad protection is particularly important.

In the present study, we designed and developed an RBD trimer as a candidate SARS-CoV-2 subunit vaccine. A single immunization elicited the acute production of protective NAbs in all *rhesus macaques*. Boost immunization induced a robust immune response, and a high titer of NAbs and a potent CD4 and CD8 T cell immune response were developed. The immune protection period lasted for at least four months (Neutralization titer > 100). Moreover, booster immunization could immediately activate the memory immune response, allowing an immunized individual to regain immune protection quickly. In addition, the vaccine-induced antibodies also had good neutralizing activity against the 501Y.V2 variant. Our vaccine candidate conferred significant protection and sterilizing immunity in vaccinated nonhuman primate lung tissue against SARS-CoV-2 infection.

## Materials and Methods

### Ethical statement

The protocols of all animal experiments were approved by Kunming National High-level Biosafety Research Center for Non-human Primates, Center for Biosafety Mega-Science, Kunming Institute of Zoology, Chinese Academy of Sciences (CAS) (permission number: IACUC20026) and performed according to the Animal Experimentation Guidelines. All applicable institutional and/or national guidelines for the care and use of animals were followed.

### Animals

Five Chinese-origin male rhesus macaques (*Macaca mulatta*, animal numbers: #140829, #140271, #163957, # 17361, #17321) weighing 4.1 to 8.7 kg and aged 3 to 6 years old were obtained from the Institute of Beijing Xieerxin Biology Resource Co., Ltd. The animals were in acceptable physical condition and were free of clinical signs of any infection. During the study period, the animals were housed individually and monitored daily for signs of illness and distress.

### Cell lines and virus

African green monkey kidney epithelial cells (Vero-E6) (ATCC, no. 1586) were used for SARS-CoV-2 virus propagation and NAb determination. The SARS-CoV-2 strain used in this experiment was provided by Kunming Institute of Zoology, Chinese Academy of Sciences (Kunming, China). The virus NMDC accession number is NMDCN0000HUI; the isolation strain is 20SF107, which was isolated and titrated in Vero-E6 cells.

### Gene construction and protein generation

The RBD region (residues Ser_325_ – Ser_530_) of the SARS-CoV-2 S protein (accession No. MN908947) was fused with an interleukin-10 signal peptide (accession No. CAG46825) at the N-terminus and foldon sequence (GYIPEAPRDGQAYVRKDGEWVLLSTFL) at the C-terminus to trimerize the protein. A C-terminal 6×His tag was also added to facilitate purification. The designed gene was synthesized by GenScript (Nanjing, China) using a mammalian codon-optimized sequence for enhanced expression and then cloned into the mammalian expression vector pCDNA3.4 (Thermo Fisher, USA) under the control of the CMV promoter. The fused protein was expressed in transfected HEK293 cells and secreted into the cell culture medium. After 72 h of culture, supernatants were harvested and purified by Ni-NTA affinity chromatography and gel filtration (Superdex 200, GE Healthcare, USA) and eluted in PBS (pH 7.2). Purified proteins were then analyzed by SDS-PAGE and Western blotting under both nonreducing and reducing conditions.

### Immunization

For the vaccination group, three rhesus macaques were vaccinated three times by the i.m. route in the deltoid muscle with a total volume of 1 ml of vaccine formulation containing 50 μg of RBD-trimer admixed with AddaVax adjuvant (InvivoGen) at weeks 0, 3, and 21, as scheduled. Blood was collected from the femoral vein into 5 ml venoject II vacuum blood collection tubes containing citric acid to evaluate the immune response. For the sham vaccination group, two macaques were vaccinated i.m. with equal amounts of PBS replacing the candidate vaccine. Serum was obtained by centrifugation at 1,200 × g at 4°C for 30 min, incubated at 56°C for 30 min, and inactivated by UV irradiation at 254 nm for 10 min before use.

### Challenge

Infectious work was performed in the Animal Biosafety Level 3 (ABSL-3) laboratory at the Kunming National High-Level Biosafety Research Center for Non-Human Primates, Kunming Institute of Zoology, Chinese Academy of Sciences (Kunming, China). Five rhesus macaques were intratracheally (60%) and intranasally (40%; 20% per naris) challenged with 1×10^7^ TCID_50_ of the virulent COVID-19 20SF107 strain 9 days after the final immunization. Rhesus macaques were monitored daily after the challenge, and their health conditions, including appetite, water drinking, behavior, appearance, and feces appearance, were recorded. When macaques showed severe clinical symptoms and became moribund, they were promptly euthanized.

### Necropsy and macroscopic pathology

Animals were euthanized one week after the challenge, and necropsies were performed in the ABSL-3 facility. Macroscopic features of necropsied tissues of the macaques were observed at dissection. For histopathology and immunohistochemistry analyses, nasal mucosa, trachea, lung, liver, spleen and kidney tissue samples collected from each animal were fixed by immersion in 4% neutral-buffered formalin for at least 7 days to kill all pathogens.

### Viral load measurement (qRT-PCR)

Viral loads in swab samples or lung tissue were determined by qRT-PCR. Viral RNA was extracted from swab samples or homogenized tissues using a High Pure Viral RNA Kit (Roche, Germany). Reverse transcription and TaqMan quantitative PCR were performed using a probe one-step real-time quantitative PCR kit (TOYOBO, Japan) on an ABI 7500 Real-Time PCR System. Primers and probes were specific for the gene encoding SARS-CoV-2 N, as follows: F: 5’-GGGGAACTTCTCCTGCTAGAAT-3’, R: 5’-CAGACATTTTGCTCTCAAGCTG-3’; probe: 5’-FAM-TTGCTGCTGCTTGACAGATT-TAMRA-3’). The 20 μl reaction mixtures were set up with 8 μl of total RNA. The cycling conditions were as follows: 50°C for 15 min, 95°C for 10 s, and 40 cycles of 95°C for 5 s and 60°C for 30 s. The dilution of each test run refers to the SARS-CoV-2 RNA Reference material (National Institute of Metrology, China). Finally, the copy number of each sample was calculated. Serial dilutions of RNA standards were included with each qRT-PCR to create a standard curve. Viral loads were expressed for the RNA Log dilutions as viral copies/g or copies/ml after calculation with the standard curve. The limit of detection was 100 copies/ml.

### ELISA

A SARS-CoV-2-RBD-specific ELISA was used to determine endpoint binding antibody titers of vaccine immune serum. Endpoint titers were defined as the reciprocal serum dilution that yielded an OD450 > 2-fold over background values. Briefly, 96-well plates were coated with 10 μg/ml recombinant RBD protein in carbonate-bicarbonate buffer (pH 9.6) at 4°C overnight. The plates were then blocked with 5% skim milk in PBS (pH 7.4) at 37°C for 1 h. Serum specimens were added to the top row (1:100), and 2-fold serial dilutions were tested in the remaining rows. The plates were incubated at 37°C for 1 h, followed by five washes with PBST. Subsequently, the plates were incubated with an HRP-conjugated anti-human IgG antibody working solution at 37°C for 30 min and washed with PBST five times. The assay was developed using 3,3’,5’,5-tetramethylbenzidine HRP substrate (TMB) with 100 μl in each well and stopped by the addition of 50 μl of 2 M H2SO4 for 10 min. Wells were measured at 450 nm by a microplate reader using Softmax Pro 6.0 software (Molecular Devices, CA, USA). All ELISA measurements were repeated at least three times with each sample in triplicate.

### Western blotting

Purified recombinant RBD protein was first separated by SDS-PAGE and then electrotransferred to a polyvinylidene difluoride (PVDF) membrane using a semidry blotting apparatus (15 V, 40 min). After transfer, the membranes were blocked with 5% skim milk in TBS buffer containing 0.1% Tween-20 for 2 h at 37°C and rinsed once with PBST. Afterward, the membranes were incubated with primary antibody (rabbit polyclonal antibodies against SARS-CoV-2-S1 at 0.1 μg/ml, prepared in blocking buffer; SinoBiological, Beijing, China). After 1 h of incubation, the membranes were washed five times with TBS containing 0.1% Tween-20, incubated with horseradish peroxidase (HRP)-conjugated goat anti-rabbit IgG (0.1 μg/ml prepared in blocking buffer) as the secondary antibody for one h, and then washed 5 times. Finally, the membranes were detected by an ECL system (CLINX, China).

### Neutralization assays

#### a) SARS-CoV-2 surrogate virus neutralization test

A surrogate virus neutralization test (sVNT) based on antibody-mediated blockage of ACE2 binding to SARS-CoV-2-RBD was performed on serum specimens using a commercially available SARS-CoV-2 sVNT Kit (L00847; GenScript, Nanjing, China) according to the manufacturer’s instructions. The serum specimens and positive or negative controls were diluted with HRP-RBD solution at a volume ratio of 1:1, i.e., 60 μl of sample was mixed with 60 μl of HRP-RBD solution. The mixtures were incubated at 37°C for 30 min. Then, 100 μl each of the mixtures was added to the corresponding 96-well plate well. The plate was covered with a Plate Sealer and incubated at 37°C for 15 min. The plate was washed four times with 260 μl of Wash Solution. Then, 100 μl of TMB solution was added to each well, and the plate was incubated in the dark at 20 - 25°C for 15 min. Finally, 50 μL of Stop Solution was added to each well to quench the reaction. The absorbance was immediately read in a microtiter plate reader at 450 nm. All measurements were repeated at least three times with each sample in triplicate.

#### b) Plaque reduction neutralization test

Live SARS-CoV-2 NAb levels were measured with a standard plaque reduction neutralization test (PRNT) in a BSL-3 facility. Briefly, serum specimens were heat-inactivated for 30 min at 56°C, serially diluted 2-fold from 1:25 to 1:12800 with DMEM containing 2% FBS in 96-well microplates, mixed with 50 plaque-forming units (PFU) of the SARS-CoV-2 20SF107 virus strain and incubated for 1 h at 37°C. The virus-serum mixtures were transferred to 12-well plates containing confluent Vero E6 cell monolayers at 37°C and with 5% CO_2_ for 1 h, and then, the virus/serum-containing medium was removed, followed by washing with PBST. Then, DMEM containing 2% FBS and 0.5% agarose was added to each well to overlay the cells and incubated for 3 days at 37°C. The cells were fixed with 10% formalin and stained with 0.2% crystal violet. Plates were washed with ultrapure water and air-dried, followed by plaque counting. Each sample was assessed in triplicate. The neutralization titer was determined by calculating the reciprocal of the highest dilution ratio of serum (neutralization endpoint) that protected 50% of cells from infection, which were defined as 50% effective concentration (EC_50_).

#### c) Pseudovirus neutralization test

A lentiviral vector pseudoneutralization test (LVV-PsN) based on antibody-mediated blockage of pseudovirus infection of cells containing SARS-CoV-2 receptors was performed on serum specimens using two commercially available SARS-CoV-2 LVV-PsN Kit (SC2087A; SC2087L; GenScript, Nanjing, China) according to the manufacturer’s instructions. Pseudoviruses bearing the SARS-CoV-2-S protein from the original Wuhan-Hu-1 strain (WT) or the 501Y.V2 (B.1.351) variant were preincubated with serially diluted positive control or serum specimens for 1 h at room temperature, and then virus-antibody mixtures (50 μl) were added to Opti-HEK293/hACE2 cells (50 μl) in a 96-well plate. After 24 h of incubation, another 50 μl of fresh DMEM was added to each well of the 96-well plate. The culture medium was discarded 48 h later, and then the cells were incubated with fresh luciferase assay substrate (Promega) for 3~5 minutes at room temperature. The fluorescence signal was detected by a multimode plate reader. Luminescence readout data were normalized to those derived from cells infected with SARS-CoV-2 pseudoviruses. The neutralization titer was determined by reducing the relative luciferase units (RLU) compared with negative control. The half-maximal inhibitory concentration (NT_50_) is calculated using nonlinear regression for the tested samples. Before experiments, serum specimens were incubated at 56°C for 30 min to inactivate the complement.

### ELISPOT

SARS-CoV-2-specific T cell responses in rhesus macaque PBMCs were assessed by an IFN-γ ELISPOT assay using recombinant SARS-CoV-2 S1 protein (GenScript, Nanjing, China). Ninety-six-well plates (Millipore) were coated with 100 μl of 10 μg/ml anti-human IFN-γ antibody (BD Biosciences, USA) per well overnight at 4°C, blocked for 1 h with sterile PBST containing 5% FBS at 37°C, and washed three times with sterile PBST. PBMCs were resuspended to 2 × 10^5^ cells/ml in RPMI 1640 medium with 10% heat-inactivated FBS (Invitrogen) and then added to the plate and subsequently stimulated with S1 protein (2 μg/ml in each well), concanavalin A (10 mM) as the positive control, and RPMI 1640 medium as the negative control. Following 20 h of incubation at 37°C, the plates were washed five times with sterile PBST and distilled water. The plates were then incubated with 2 μg/ml biotinylated anti-human IFNγ antibody for 1 h at room temperature, washed six times with sterile PBST, and incubated for 1 h at room temperature with streptavidin–HRP. After five washes with sterile PBST and one with sterile PBS, AEC substrate solution (Abcam) was added to the plates. When the spots were clear enough, the reaction was stopped by washing with distilled water and air-dried, and spots were counted with an ELISPOT Analysis System (At-Spot-2100, China). Assays of all specimens were performed in triplicate. Cells alone in the absence of stimulants were used as a negative control. The numbers of spot-forming cells (SFCs) per 10^6^ cells were calculated. The medium background levels were typically < 15 SFC per 10^6^ cells.

### ICS and flow cytometry

SARS-CoV-2-specific CD4^+^ and CD8^+^ T cell responses in NHP PBMCs were assessed by flow cytometry using recombinant SARS-CoV-2 S1 protein (GenScript, Nanjing, China). Briefly, NHP PBMCs were added to a plate (1×10^6^/well) and then stimulated with S1 protein (2 μg/ml in each well) or RPMI 1640 medium as a control for 5 h. Subsequently, the cells were incubated with GolgiPlug/GolgiStop (BD Biosciences, USA) to block intracellular cytokine transport for 6 h at 37°C. Afterward, the cells were washed and surface stained with antibodies against human CD4 (FITC-conjugated), CD8α (FITC-conjugated), and CD44 (FITC-conjugated) for 30 min at room temperature. Subsequently, cells were fixed and permeabilized with a Cytofix/Cytoperm kit (BD Biosciences, USA). Cells were then washed and incubated with anti-human IFN-γ antibodies (PE-conjugated) for 1 h at room temperature, washed with permeabilization buffer, and fixed in 1% paraformaldehyde. All specimens were processed with a FACSCanton flow cytometer (BD Biosciences, USA).

### Histopathological analysis

Lung tissues were fixed overnight in 4% paraformaldehyde (PFA), dehydrated, and embedded in paraffin. Subsequently, 5-μm-thick tissue sections were processed for histology by hematoxylin and eosin staining as previously described. The immunohistochemical reaction was performed on paraffin-embedded lung sections using a rabbit anti-SARS-CoV-2-NP monoclonal antibody followed by an HRP-conjugated goat anti-rabbit secondary antibody as described. After the DAB chromogenic reaction, nuclear counterstaining, dehydration and mounting, tissue staining was visualized under a microscope, and images were acquired and analyzed.

### Statistical analysis

All statistical analyses were performed using GraphPad Prism 8.0 software (GraphPad Software, CA, USA). Statistical significance among different groups was determined by Student’s t-test, and differences between the experimental groups were considered statistically significant at the P<0.05 level.

## Results

### Construction of a SARS-CoV-2 RBD trimer vaccine

To simulate the native SARS-CoV-2 S protein trimeric form and improve conformational homogeneity, a natural trimerization domain of T4 bacteriophage fibritin (foldon) was fused to the C-terminus of the SARS-CoV-2 RBD protein (Fig. 1A). An interleukin-10 signal peptide was added to the N-terminus of the RBD to improve peptide secretion. The recombinant protein, named RBD-trimer, was ectopically expressed in human embryonic kidney (HEK293) cells. RBD-trimer was purified by Ni-NTA affinity chromatography and gel filtration. The purity of the recombinant protein was assessed by SDS-PAGE (Fig. 1B) and Western blot (Fig. 1C) assays with or without reducing conditions. Notably, RBD-trimer did not dissociate into monomers with heating, indicating the thermal stability of this protein complex. According to the N-terminal sequencing of RBD-trimer, the interleukin-10 signal peptide was efficiently cleaved off. We chose the AddaVax adjuvant, an MF59-like nanoemulsion, to prepare our vaccine formulation to better stimulate cellular immune responses. The AddaVax-adjuvanted RBD-trimer protein was used throughout the following immunizations in animals.

**Fig. 1.**
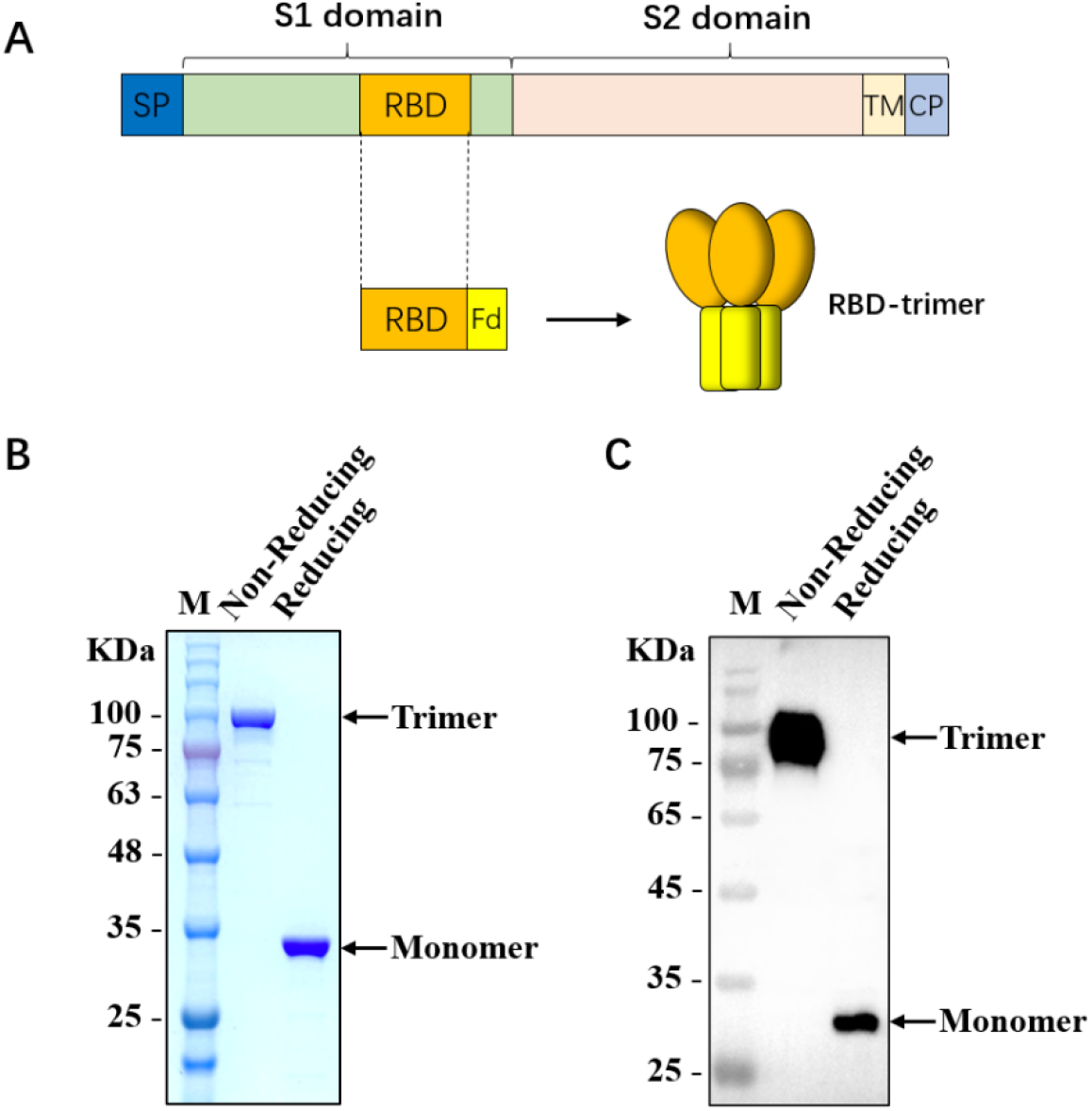
Characterization of SARS-CoV-2 RBD-trimer. (A)Schematic design of SARS-CoV-2 RBD-trimer. (B-C) The purified recombinant protein was analyzed by SDS-PAGE (B) and Western blot with a polyclonal antibody against SARS-CoV-2-S1 (C). M: Marker. 10 μg of recombinant protein was loaded in each lane.

### Immunization with RBD-trimer elicited robust humoral and cellular immune responses in *rhesus macaques*

To assess the immunogenicity of the SARS-CoV-2 RBD-trimer vaccine, five male *rhesus macaques* (3-6 years old) were vaccinated with either RBD-trimer or phosphate-buffered saline (PBS). The animals were intramuscularly immunized with 50 μg of RBD-trimer with AddaVax adjuvant (n=3, animal numbers: #140829, #140271, #163957) or PBS (n=2, animal numbers: # 17361, #17321) at 0, 3 and 21 weeks. All the macaques were phlebotomized as outlined in Fig. 2A. Serum was isolated for ELISA and neutralizing activity tests, in which NAbs were measured by a standard plaque-reduction neutralization test (PRNT), a lentiviral vector pseudoneutralization test (LVV-PsN), and a surrogate virus neutralization test (sVNT) based on antibody-mediated blockage of ACE2 binding to the RBD. The NAb average titers of the vaccinated animals reached approximately 100 (EC_50_)/200 (sVNT) and 1000 (EC_50_)/4000 (sVNT) after the first and second immunizations, respectively, and then gradually decreased to 130 (EC_50_)/400 (sVNT) at week 21. Considering a previous report that NAbs can achieve full immune protection as long as the titer reaches 50 (EC_50_) ^[5]^, these data suggest that our RBD-trimer vaccine can induce acute and durable protective immunity for no less than 4 months in macaques. The second boost produced extremely high titers of NAbs (2600 EC_50_, 8000 sVNT) (Fig. 2B, C and Fig. S1A). Similar to the NAb results, RBD-specific ELISA IgG titers in the vaccination group gradually increased and peaked after the boosts (Fig. 2D). To evaluate whether the 501Y. V2 variant is resistant to neutralization by RBD-trimer vaccinee serum, we tested the neutralizing activity against the original SARS-CoV-2 strain (WT) and 501Y.V2 variant on day 7 post-second boost using the LVV-PsN assay, and the results showed that the NAb average titer was approximately 1030 (NT_50_) against the 501Y.V2 variant, which is approximately 30% lower than that against the WT strain (NT_50_ of 1500) (Fig. 2E and Fig. S1B). By contrast, the neutralizing activities of mRNA vaccinee serum against the 501Y.V2 variant were significantly lower (41.2-fold reduction, Pfizer; 20.8-fold reduction, Moderna) ^[25, 26]^, suggesting that our vaccine can protect against infection with the 501Y.V2 variant. Neither RBD-specific binding antibodies nor NAbs were detected in the sham-vaccinated macaques.

**Fig. 2.**
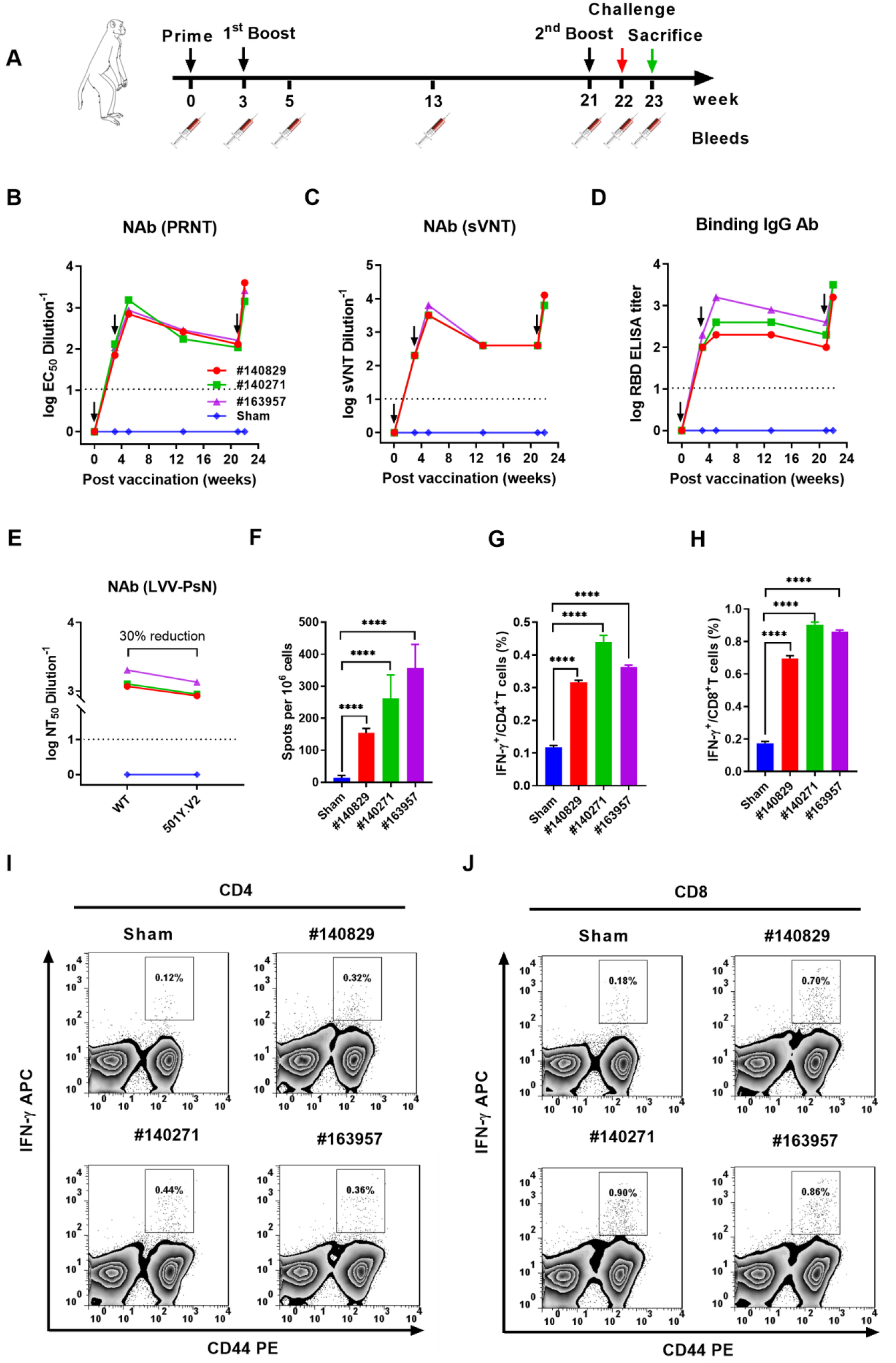
SARS-CoV-2 RBD-trimer induced robust humoral and cellular immune responses in *rhesus macaques*. Flow chart of experimental design (A). Arrows indicate weeks of immunization (black), challenge (red), and sacrifice (green), syringes indicate sampling weeks. The serum samples were used to test NAb titers by PRNT assay (B) and a surrogate virus neutralization test (sVNT) (C) and RBD-specific ELISA antibody titers (D). Neutralization of the original Wuhan-Hu-1 (WT) and 501Y.V2 variant viruses by the second-boost serum was measured with the use of a recombinant lentiviral-based pseudovirus neutralization assay (LVV-PsN) (E). Measurements below the detection limit were assigned a value of 1. PBMCs isolated from immunized and sham groups 2 weeks after the first boost were collected for IFN-γ detection by ELISPOT assay after stimulation with S1 protein (F). The proportion (G, I) of CD4^+^ T cells secreting IFN-γ among all CD4^+^ T cells and the proportion (H, J) of CD8^+^ T cells secreting IFN-γ among all CD8^+^ T cells were evaluated by intracellular cytokine staining. The dotted lines indicate the detection limit. Data are shown as the means ± SEM (standard errors of the mean). *P* values were analyzed by Student’s t-test (****, P < 0.0001).

Peripheral blood mononuclear cells (PBMCs) were isolated after the first booster immunization, and SARS-CoV-2-specific CD4+ and CD8+ peripheral blood T cell responses were measured by IFN-γ enzyme-linked immunospot (ELISPOT) and intracellular cytokine staining (ICS) assays. Compared to that of the sham group, IFN-γ secreted by PBMCs was significantly increased after SARS-CoV-2-S1 stimulation (140-440 spots/10^6^ cells from 7-21 spots/10^6^ cells, Fig. 2F). Moreover, the proportion of IFN-γ-secreting CD4^+^ T cells (IFN-γ/CD4^+^) in all CD4^+^ T cells increased from 0.12% to 0.32-0.44% (Fig. 2G, I), while the proportion of IFN-γ-secreting CD8^+^ T cells (IFN-γ/CD8^+^) reached 0.70-90% from 0.18% (Fig. 2H, J), demonstrating that immunization with RBD-trimer stimulated a potent specific cellular immune response.

### The RBD-trimer vaccine provides complete protection against SARS-CoV-2 challenge in *rhesus macaques*

We next assessed the protective efficacy of the RBD-trimer vaccine against SARS-CoV-2 infection in *R. macaques*; all the macaques were intranasally and tracheally challenged with 1 × 10^7^ TCID_50_ of the SARS-CoV-2 20SF107 strain at 9 days after the second booster vaccination (Fig. 2A). Throat, nose, and rectal swabs and tracheal brush specimens were collected over a time course post-infection. All animals were euthanized at 7 days post-infection (dpi) to assess the viral load in the respiratory tissues. A high virus burden was detected in the swab specimens from the throat, nose, and rectum and tracheal brushes throughout the control animals’ detection periods (Fig. 3 A, B, C, D). The virus loads in nasal swabs and tracheal brushes of control macaques gradually decreased after reaching peak values at 2 dpi. In comparison, the virus loads in anal swabs increased gradually from 2 dpi. In contrast, a mild viral infection was detected in the swabs and tracheal brushes at 1 dpi and then quickly dropped to an extremely low level in the RBD-trimer-immunized macaques. All seven different lung lobe tissues from the control macaques showed high levels of viral genomic RNA. In contrast, no viral RNA was detected in the immunized macaques, indicating that our RBD vaccine candidate could confer sterilizing immunity in lung tissues (Fig. 3E). Additionally, no clinical signs were found in the immunized macaques after challenge, while the control macaques exhibited symptoms, including cough and increased body temperature (Fig. 3F).

**Fig. 3.**
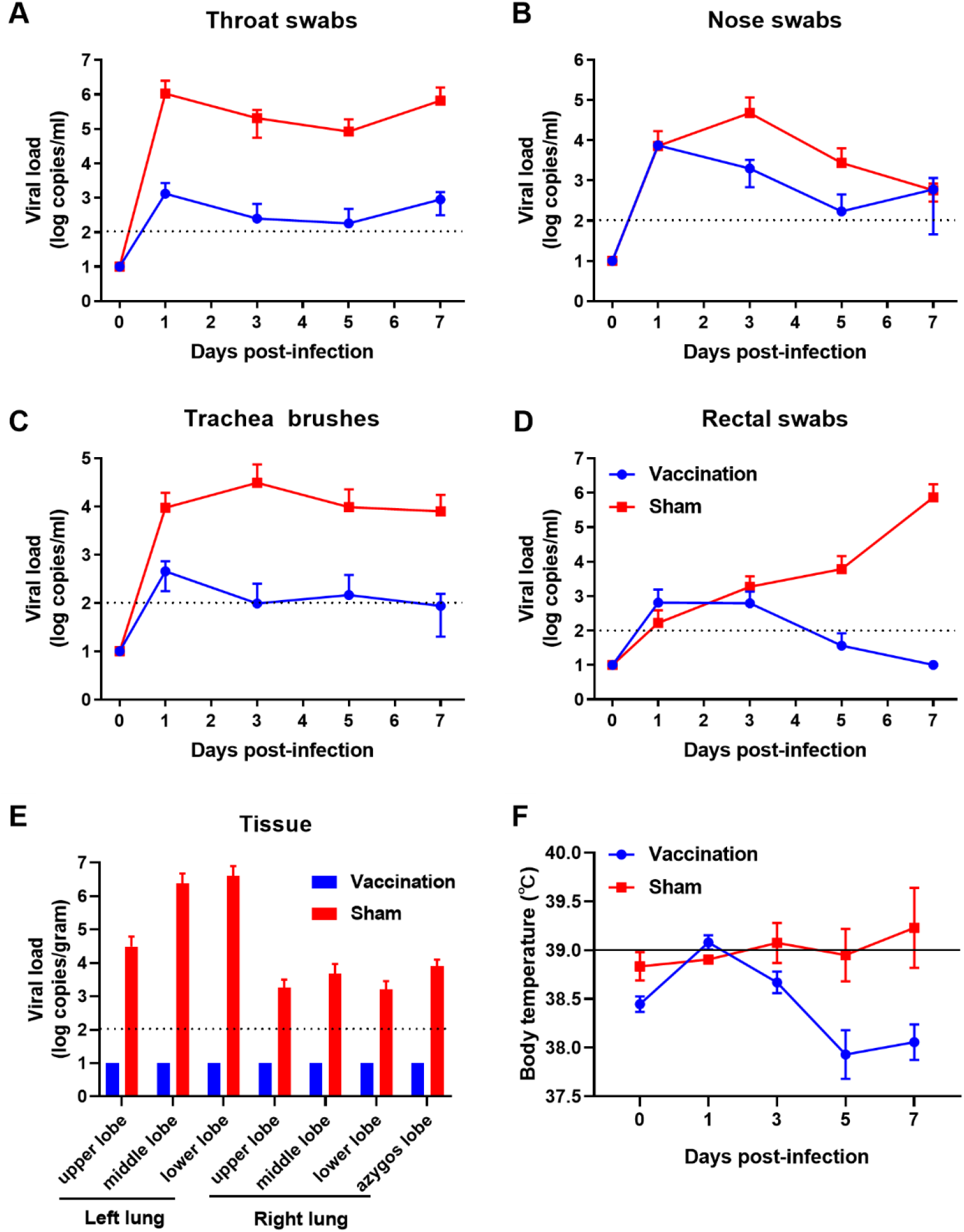
Protective efficacy of the RBD-trimer vaccine in *rhesus macaques*. Macaques were challenged intranasally and tracheally with SARS-CoV-2. Viral loads in throat swabs (A), nasal swabs (B), tracheal brushes (C), rectal swabs (D), and lung tissues (E) were measured by qRT-PCR. Changes in body temperature of the animals (F). The dotted lines indicate the detection limit. Measurements below the limit of detection were assigned a value of 10. The solid lines represent the upper limit of the standard body temperature reference value.

Histopathological analysis of the fixed lungs was performed with hematoxylin-eosin (H&E) staining. No visible pathological change or inflammatory cell infiltration was observed in the lung tissues from immunized macaques (Fig. 4A, B). In contrast, severe histopathological changes were found in the control macaques, including apparent thickened alveolar walls, mononuclear inflammatory cell infiltration, focal exudation, hemorrhage, and architecture disappearance (Fig. 4C, D). Immunohistochemical (IHC) assays indicated that multiple pneumocytes in the lung sections from the control macaques were positive for SARS-CoV-2 N protein staining (Fig. 4G, H). By contrast, no viral antigen was detected in the lung tissues from the immunized animals (Fig. 4E, F).

**Fig. 4.**
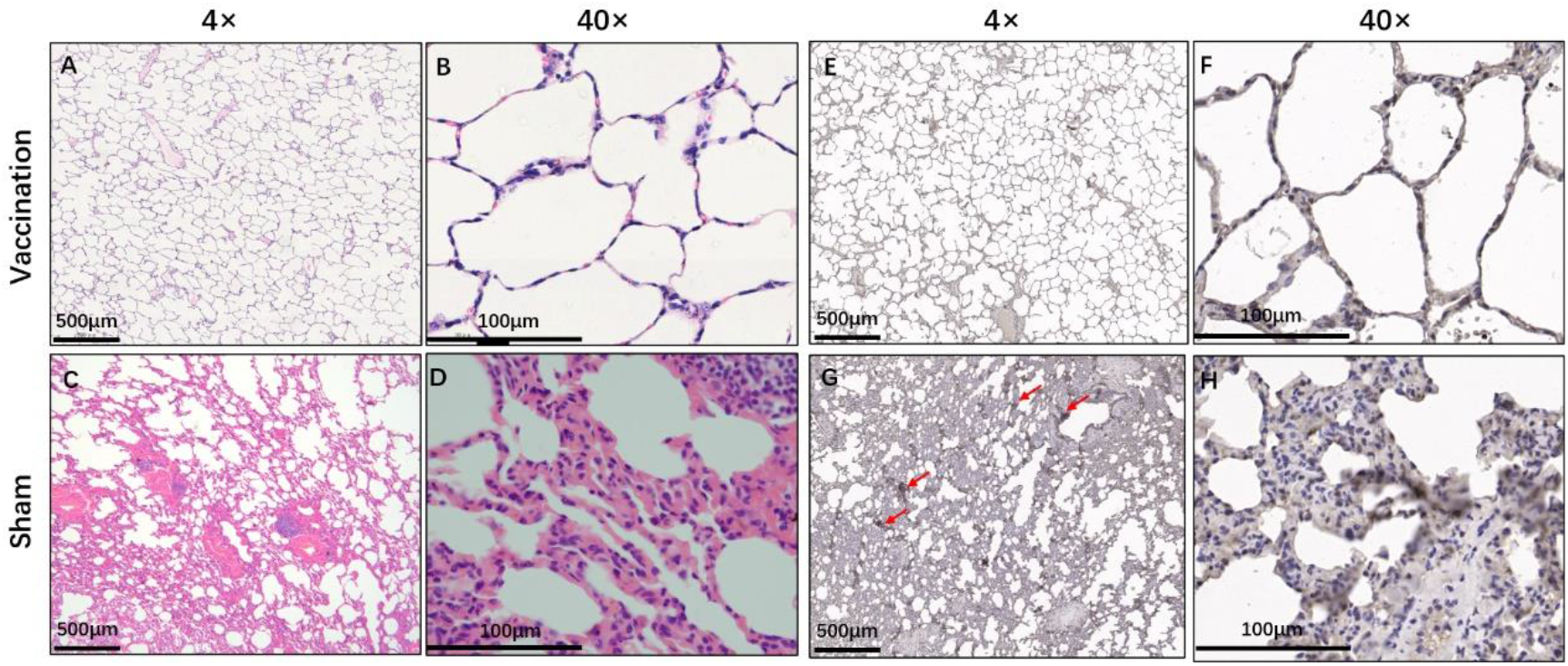
H&E staining and IHC analysis of lungs from rhesus *macaques*. Macaques immunized with RBD-trimer or PBS were sacrificed at 7 dpi. Lung tissue sections were stained by H&E staining (A, B, C, D). IHC detection of the N protein of SARS-CoV-2 (E, F, G, H). Immunohistochemically positive cells are marked by red arrows. Images of different tissue pathologies at low magnification (4×) and higher magnification (40 ×) are shown.

**Fig. 5.**
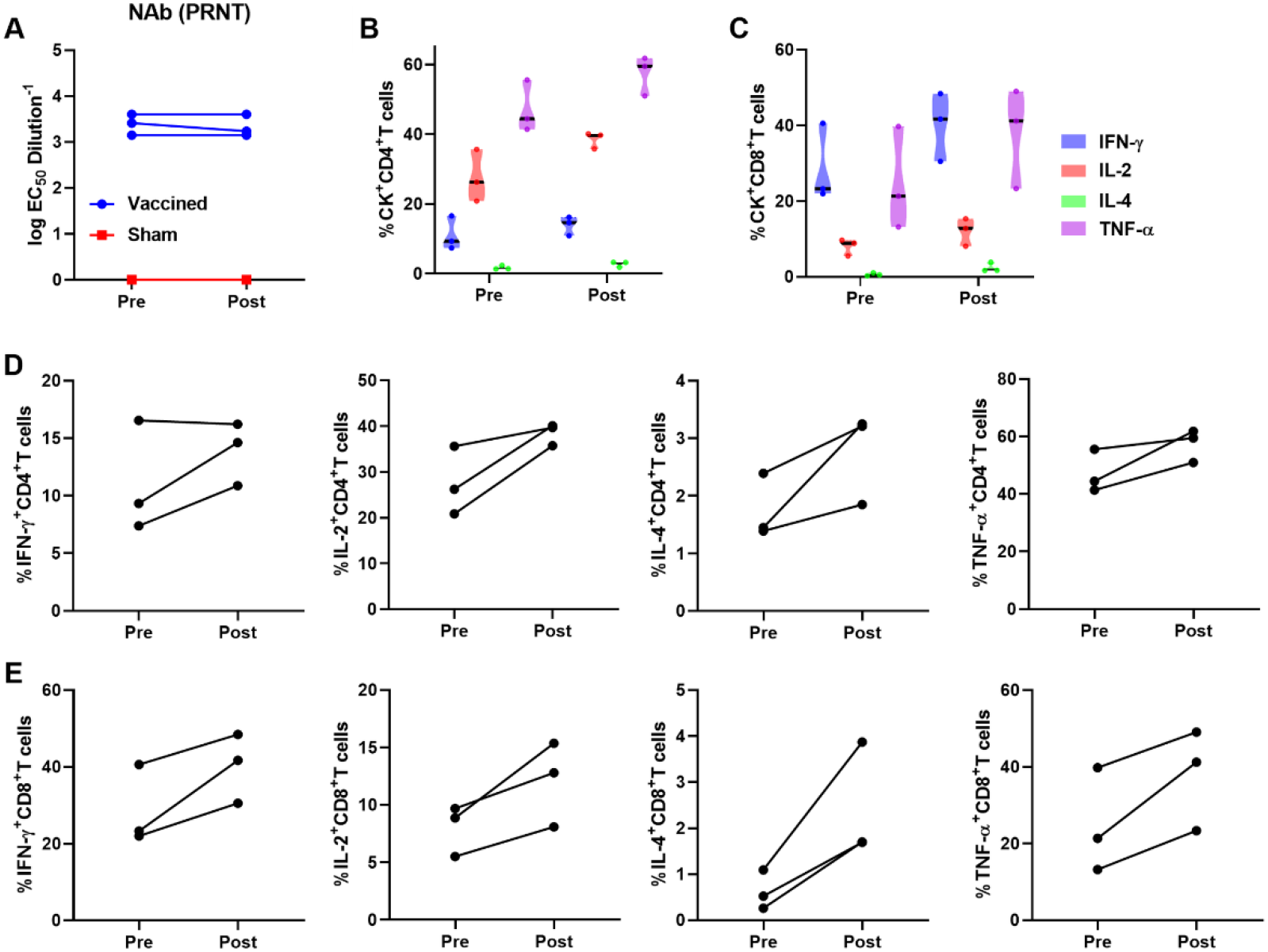
NAb and memory T cell responses induced by SARS-CoV-2 infection in vaccinated *rhesus macaques*. Serum and PBMCs were collected and isolated from macaques before infection (Pre) and 7 dpi (Post). Serum NAb titers were measured by PRNT assay (A). Cells were stained for intracellular production of IFN-γ, IL-2 IL-4 and TNF-α. The grouped violin plot graphs show the mean changes in the total percentage of CD4+ and CD8+ T cells expressing the indicated cytokine in the blood (B, C). Each line represents an individual animal (D, E).

To evaluate anamnestic humoral and T cell responses in vaccinated *rhesus macaques* after challenge, paired serum and PBMCs were collected from vaccinated animals before and after SARS-CoV-2 challenge, and changes in the NAb titer and the percentage of cytokine-expressing CD4+ and CD8+ T cells were analyzed. The results showed that the NAb titer did not change one week after viral challenge, indicating that no anamnestic humoral immune response was induced. When the vaccine candidate fails to induce a sufficient protective immune response ^[11]^, the challenge enhances NAb titers; therefore, our vaccine protection is sterilizing. Simultaneously, the proportions of CD4 and CD8 T cells secreting IFN-γ, IL-2, IL-4 and TNF-α increased significantly (approximately 1.5-fold or 2.5-fold increase in CD4 or CD8 T cells), suggesting that the SARS-CoV-2-specific memory T cell immune response was stimulated to secrete cytokines.

Hematology and serum biochemistry values of all macaques were tested over a time course post-infection. The number of platelets in the control macaques increased and reached a peak at 5 dpi, indicating hemorrhage in the tissues (Fig. S2), which suggested blood cell exudation in the lung blood vessels. In addition, the number of lymphocytes in the control macaques slightly increased. The concentrations of aspartate aminotransferase (AST) and alanine aminotransferase (ALT) in the serum of the control macaques increased and exceeded the upper reference range at 1 and 2 dpi, suggesting that these macaques had liver injury after SARS-CoV-2 challenge (Fig. S3). By contrast, the hematological and biochemical states of immunized macaques stayed at a normal range during the viral challenge periods.

Compared with young people, the frail middle-aged population has higher risks of severe COVID-19 with a higher mortality rate, and the virus replicates more efficiently in their lungs and other organs^[28]^. NHP model animals infected with SARS-CoV-2 showed that older macaques developed extreme viral replication in the lungs^[29]^. Therefore, the protective effect of vaccines for middle-aged and frail people is highly valuable. Thus, our vaccination group included a frail macaque (#140829, 6 years old, 4.8 kg), which was only half the weight of macaques of the same age. This animal can better simulate the candidate vaccine’s immune and protective efficacy for frail people. Compared to the other two macaques, this animal had a slightly weaker immune response and lower antibody levels during the first 2 vaccinations. However, after the third immunization, its immune responses were no different from those of the other macaques (Fig. 2B, C and Fig. S1A). This frail monkey was fully protected by RBD-trimer vaccination, suggesting that the vaccine could provide adequate protection to middle-aged and frail people against COVID-19.

## Discussion

The severe COVID-19 pandemic and the precipitously increasing numbers of deaths worldwide necessitate the urgent development of SARS-CoV-2 vaccines. Global researchers are racing toward safe and effective vaccines against COVID-19 based on multiple platforms, and some vaccines have received emergency use validation, including mRNA vaccines (Moderna, Pfizer/BioNTech), inactivated virus vaccines (Sinopharm, Sinovac, Janssen), and adenovirus vector-based vaccines (AstraZeneca, CanSino). In this study, an engineered trimeric RBD subunit vaccine candidate elicited robust humoral and cellular immune responses in nonhuman primates (NHPs), resulting in significant protection against SARS-CoV-2 infection.

To improve the immunogenicity of the recombinant RBD protein and better mimic the native trimeric SARS-CoV-2 S structure, we fused a trimer motif to make the RBD form a stable trimer structure; the construct will not dissociate even if the temperature is above 60°C, and there was no change after 3 months of storage at 4°C. Western blot analysis showed that compared with the monomer, the trimer binds to the SARS-CoV-2 S1 polyclonal antibody more strongly, suggesting that the conformational epitope is better retained. In contrast with the results of other vaccine candidates reported in the NHP model ^[5]^, NHPs can obtain protective NAb titers (EC_50_ titer of 100) with a single dose of our candidate, suggesting that it can be used for emergency vaccination of high-risk populations. There are few reports about the persistence of NAb responses induced by COVID-19 vaccine candidates. To evaluate the candidate’s protection persistence, our immunity cycle spanned half a year. The results showed that the antibody level gradually decreased to an EC_50_ average titer of 50 4 months after the second vaccination. Although the antibody level was much lower than that immediately after vaccination, it was still above the protective antibody level, suggesting that this vaccine can achieve no less than 4 months of long-lasting humoral protection. Considering that SARS-CoV-2 may coexist with humans for a long time, a vaccine that can generate acute and durable immune protection is of great value. Due to the high variability of SARS-CoV-2, multiple variants have recently been identified and are now spreading worldwide, such as B.1.1.7 (501Y.V1), B.1.351 (N501Y.V2) and P1 (N501Y.V3), which appear to be more easily transmitted ^[30]^. There is a growing concern that the N501Y.V2 and N501Y.V3 variants could impair the protective efficacy of licensed vaccines ^[26]^. Therefore, the broad-spectrum protection of the vaccine is also a focus of our attention. Previous research reported that the AddaVax adjuvant, a squalene-based oil-in-water emulsion similar to MF59, could increase the broad protection of the influenza virus vaccine by stimulating a cross-protective humoral response ^[31]^. Compared with aluminum adjuvants, squalene-based adjuvants have been shown to induce humoral and cellular immune responses in a more balanced manner ^[32]^. Therefore, we chose this adjuvant to formulate our candidate vaccine. Excitingly, the antibodies produced in response to our vaccine showed good neutralizing activity against N501Y.V2 variant.

NAbs are considered to play a vital role in preventing SARS-CoV-2 infections. Monoclonal antibodies (MAbs) and convalescent plasma have been used to treat severe COVID-19 and have shown sound therapeutic effects. A MAb regimen has been authorized by the US Food and Drug Administration to treat COVID-19. Therefore, the NAb titer induced by a vaccine is a crucial parameter for vaccine development. The cellular immune response is also a critical effector in virus clearance and alleviation of clinical symptoms. Moreover, studies have shown that the duration of the memory T cell response induced by infection is much longer than that of antibody levels, so cellular immunity plays a crucial role in the durability of vaccine protection ^[33]^. In addition, as the level of NAbs gradually decreases after vaccination, nonneutralizing antibodies may promote infection via the ADE effect, but virus-specific T cell immunity can reduce this potential risk. An ideal vaccine should be able to induce humoral and cellular immune responses in a balanced manner. Compared with the RBD monomer, RBD-trimer generated a higher level of NAbs, resulting in more potent protection. In addition, it is surprising that by formulating an Addavax adjuvant, our vaccine candidate also demonstrated a strong CD4+ and CD8+ T cell immune response in NHPs, suggesting more robust protection.

SARS-CoV-2 infects hosts mainly in the respiratory tract and lung, and pneumonia is the typical symptom of COVID-19; our data correspond to this report. Sham-immunized macaques showed severe pneumonia; in contrast, macaques immunized with our vaccine candidate exhibited significant protection of the lung with viral clearance. The main changes in sham-immunized macaques were inflammatory granulocyte infiltration, disseminated thickening of alveolar septa, and hemorrhage. By contrast, immunized macaques showed less severe lung symptoms. Hematological analysis showed that the platelet concentration of the sham-immunized macaques continued to increase and reached a peak at 5 dpi, suggesting visceral bleeding. In contrast, the immunized macaques did not have any hematological abnormalities. Furthermore, no viral RNA was detected in the lungs of the immunized macaques by qPCR or immunohistochemical assays. By contrast, in the lungs of sham-immunized macaques, the viral RNA level reached almost 10^7^ copies/gram. These results proved that our vaccine candidate conferred sterilizing immunity in the lung tissue of immunized NHPs. SARS-CoV-2 attacks host organs beyond the lung; liver injury has also been recorded. In this work, after challenge, the ALT and AST values of the unimmunized animals increased, indicating that SARS-CoV-2 attacked their liver. There were no abnormalities in the vaccinated animals, which means that our vaccine candidate may protect the animals from liver injury. In short, our candidate vaccine can effectively protect the organs of NHPs from virus attacks.

Since the antibody level induced by infection or vaccination cannot last overly long, when the antibody level of the population drops to a level insufficient to achieve immune protection, there is a risk of reinfection. Once there is another outbreak, how can individuals quickly obtain immune protection? Our results proved that only one boost could quickly regain complete immune protection for the vaccinated NHPs. Notably, after the COVID-19 pandemic is under control, if SARS-CoV-2 infection appears again, people who have been vaccinated can be immediately administered a booster immunization to avoid another large-scale outbreak of the epidemic.

Multiple mammalian species, including rats, mice, guinea pigs, rabbits, cynomolgus macaques, and rhesus macaques, have been used for COVID-19 animal models. It is still too early to define the most suitable animal model for studying SARS-CoV-2 infections. Because of their close genetic relation to humans, NHPs are usually used as animal models to evaluate vaccines and drugs against human disease. We provided evidence for trimeric RBD subunit vaccine safety in NHPs and did not observe infection enhancement or immunopathological exacerbation in our studies. Our data also demonstrate significant protection against SARS-CoV-2 challenge with 50 μg per vaccination dose in NHPs.

In summary, our RBD-based subunit vaccine candidate induced robust humoral and cellular immune responses and protected NHPs against SARS-CoV-2 infection, and it also showed perfect safety. There were no abnormalities in body temperature, weight, clinical signs, pathology, hematology, or biochemical indicators during vaccination and challenge. These data proved the possibility that our trimeric RBD can be a promising candidate vaccine to prevent COVID-19.

## Acknowledgments

We thank JFS and JHZ from Beijing Primabio Inc for their assistance with husbandry and experiments. This study was funded by National Key Technologies Research and Development Program grants 2018YFC1200600 and 2018YFC1200500 to LMY. and a grant from Strategic Priority Research Program of Chinese Academy of Sciences XDB29010000 to WJL.

## Author contributions

Conceptualization: LMY, WJL; Methodology: LMY, DYT, JBH, WHF; Investigation: LMY, YZ, WQS, YLL, XDT, YQW, DDY, XLF; Visualization: LMY, YHB; Funding acquisition: LMY, WJL; Project administration: WJL, LMY; Supervision: WJL, YTZ; Writing – original draft: LMY, DYT; Writing – review & editing: WJL, YHB, GC

---

**Fig. S1.**
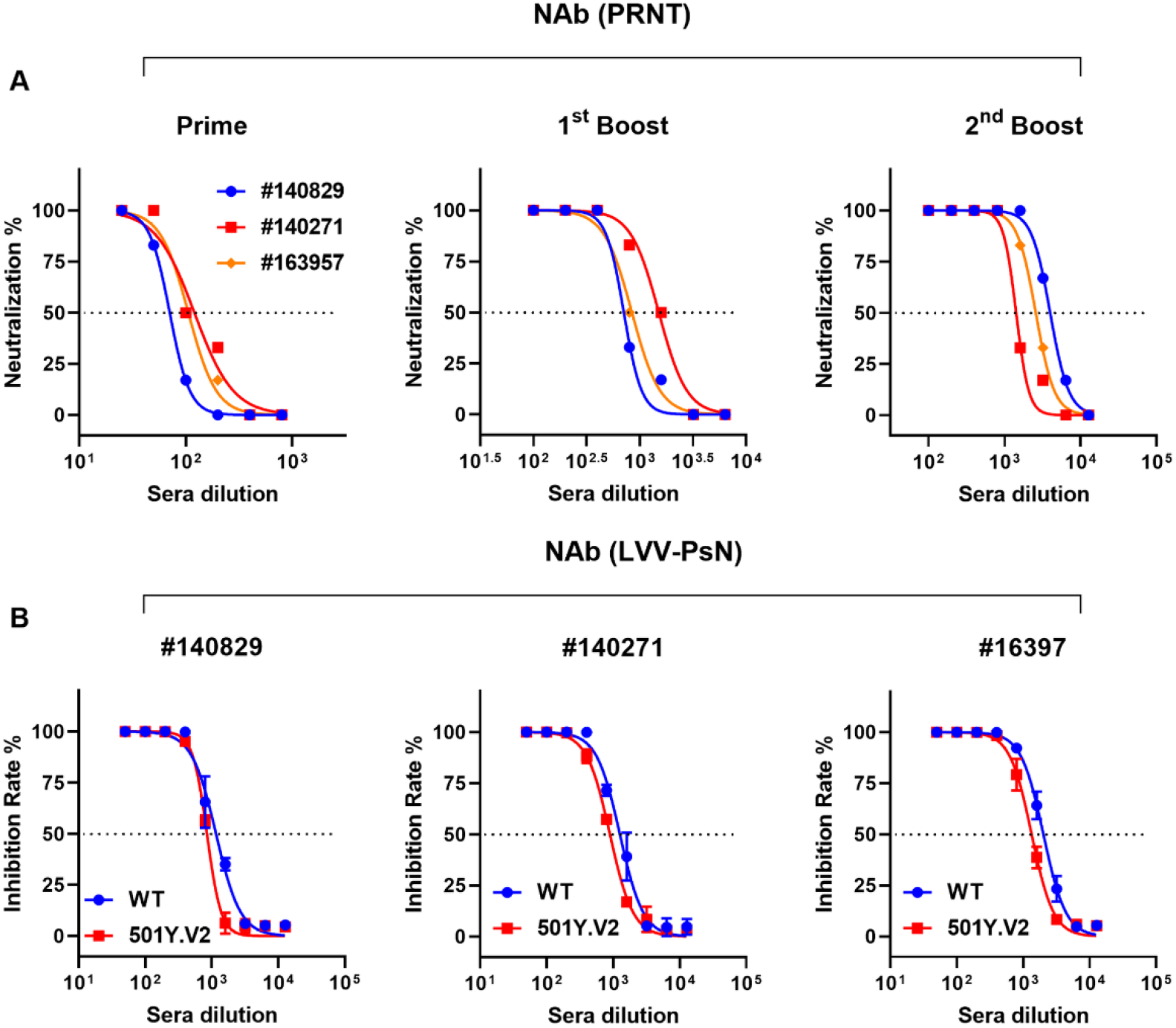
*In vitro* neutralization assays of immunized *rhesus macaque* serum. The serum samples after three immunizations were used for testing NAb titers by PRNT assay (A), respectively. The sera samples collected on day 7 post the 2^nd^ boost were used for testing NAb titers by LVV-PsN assay (B). Serial dilutions of serum were pre-incubated with Pseudoviruses bearing the SARS-CoV-2-S protein from the original Wuhan-Hu-1 strain (WT) or the 501Y.V2 (B.1.351) variant, and the percentage inhibition of cell infection were determined. Nonlinear regression was performed using a log (inhibitor) versus normalized response curve and a variable slope model (R2> 0.95 for all curves). The EC_50_ (PRNT) and NT_50_ (LVV-PsN) were calculated by using the Reed-Muench method. Data are shown as means ± SEM.

**Fig. S2.**
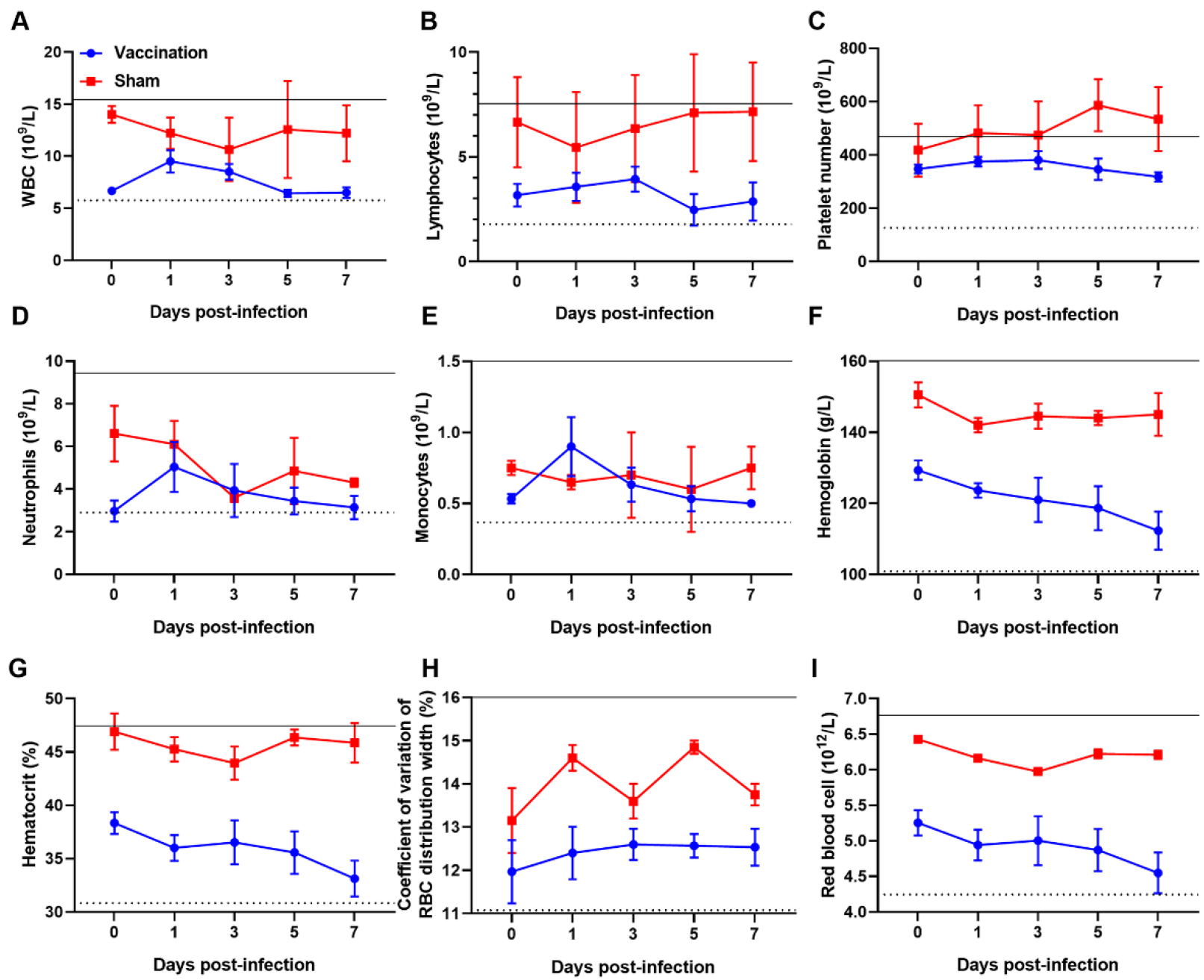
Hematological index changes of rhesus macaques after SARS-CoV-2 infection. Macaques were challenged, and hematological values were obtained before and 1, 3, 5, and 7 days after challenge. The solid lines and the dotted lines represent the upper and lower limits of the normal reference range.

**Fig. S3.**
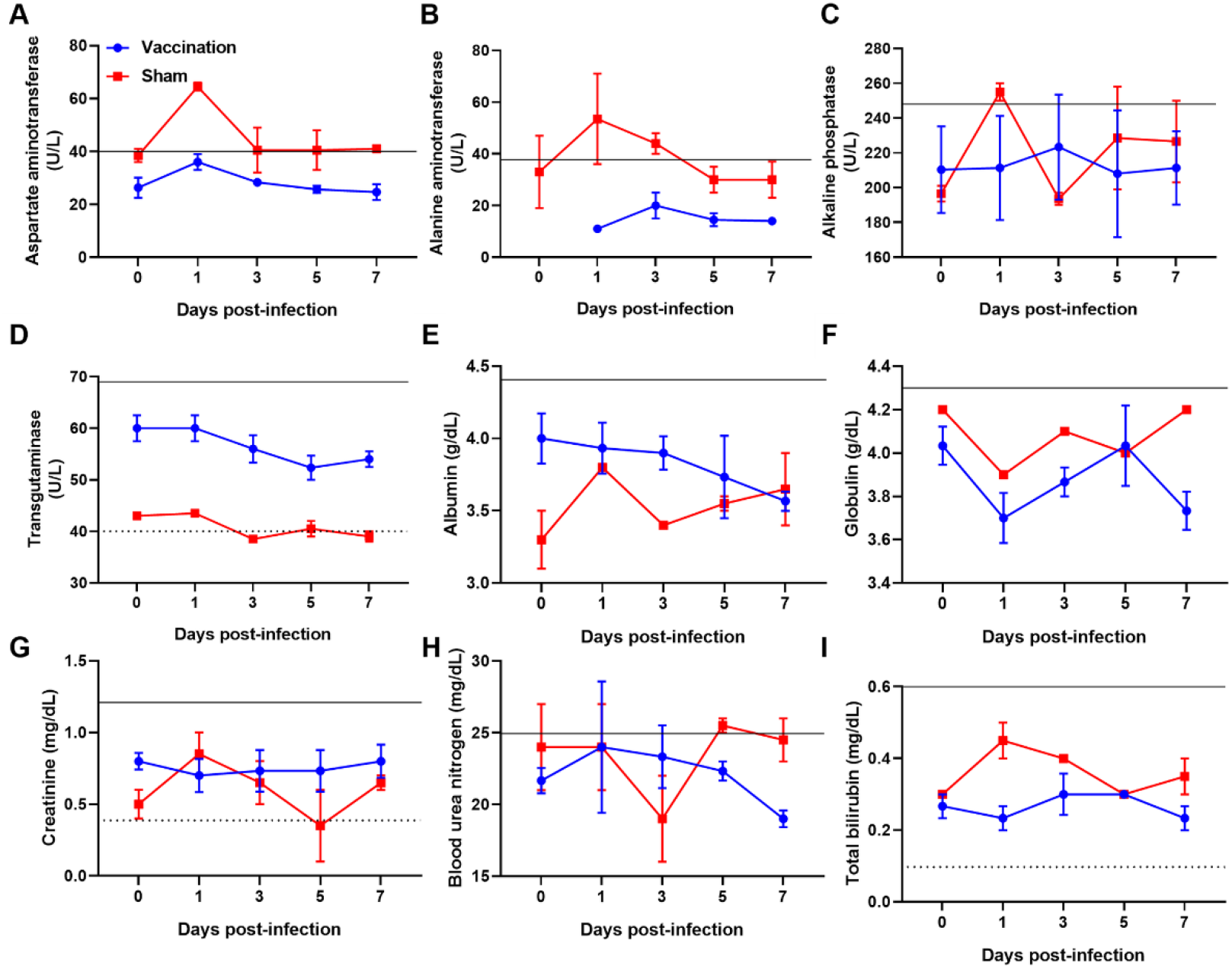
Serum biochemical index changes of rhesus macaques after SARS-CoV-2 infection. Macaques were challenged, and body temperature and serum biochemical values were measured before and 1, 3, 5, and 7 days after the challenge. The solid lines and the dotted lines represent the upper and lower limits of the normal reference range.

